# The role of diffusion MRI in neuroscience

**DOI:** 10.1101/140459

**Authors:** Yaniv Assaf, Heidi Johansen-Berg, Michel Thiebaut de Schotten

## Abstract

Diffusion weighted imaging has further pushed the boundaries of neuroscience by allowing us to peer farther into the white matter microstructure of the living human brain. By doing so, it has provided answers to fundamental neuroscientific questions, launching a new field of research that had been largely inaccessible. We will briefly summarise key questions, that have historically been raised in neuroscience, concerning the brain’s white matter. We will then expand on the benefits of diffusion weighted imaging and its contribution to the fields of brain anatomy, functional models and plasticity. In doing so, this review will highlight the invaluable contribution of diffusion weighted imaging in neuroscience, present its limitations and put forth new challenges for the future generations who may wish to exploit this powerful technology to gain novel insights.

---

“*We admire the contrivance of the fibre of every muscle, and ought still more to admire their disposition in the Brain, where an infinite number of them contained in a very small space, each execute their particular offices without confusion or disorder*” ^1^.

As noted by Steno, post mortem dissections of the brain reveal an astonishing level of complexity, particularly within the white matter fibre pathways, composed of trillions of axons. As Isaac Newton intuitively suggested a few years later, these axons propagate electricity “*along the solid filament of the nerves, from the outward organs of sense to the brain, and from the brain into the muscles*” ^2^. **Anatomical exploration** of the white matter connections of the human brain then becomes a new challenge, mixing medical knowledge with advanced practical skills of dissection and illustration (**Figure 1**).

**Figure 1:**
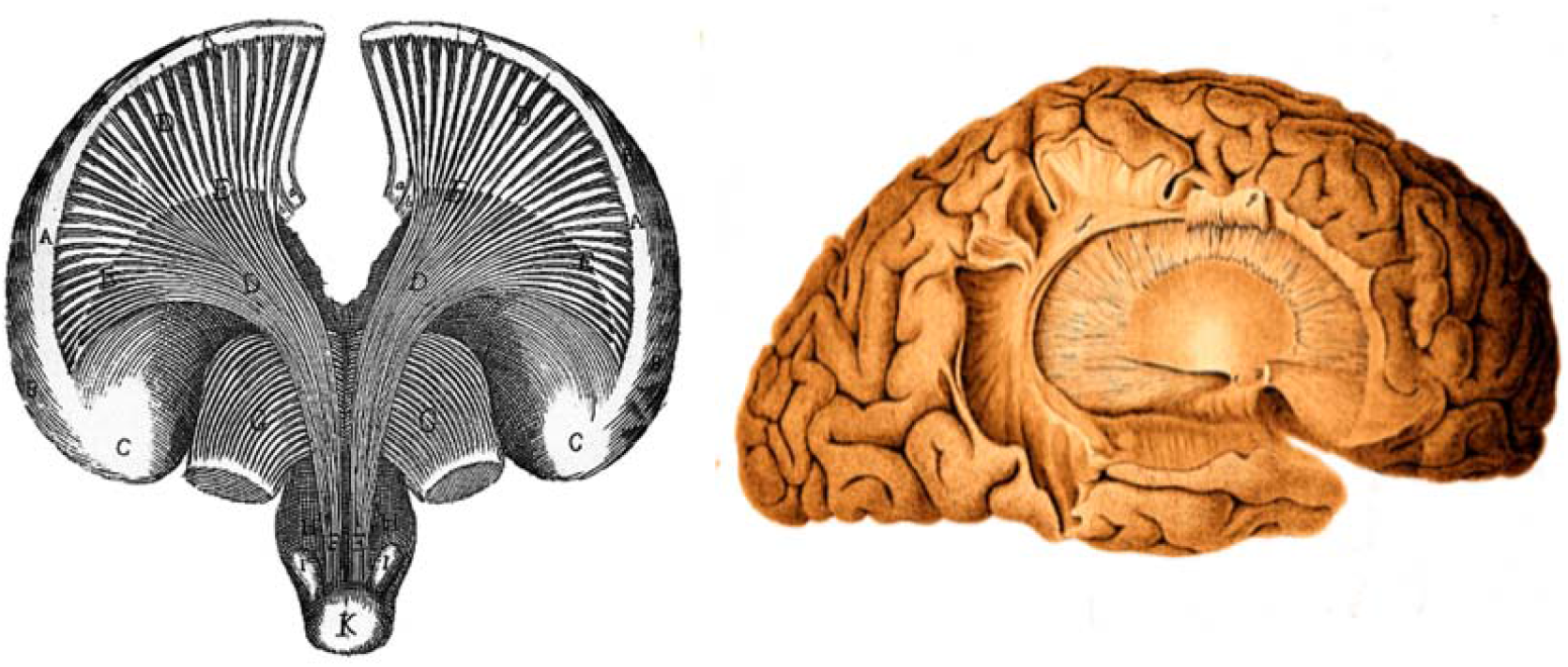
First drawings of white matter connection in the human brain. Left, brain dissection from Vieussens ^3^ and right, brain dissection from Johann Christian Reil ^4^.

These anatomical descriptions were a stepping-stone for the elaboration of new **theories of brain function**. This can be seen for example toward the end of the XIXth century, when Meynert extended Newton’s vision even further, by introducing the possibility of reasoning occurring through the association of specialised areas of the brain via their white matter connections. The concept of associationism thus was born ^5^.

Empirical validation of associationism came soon after with the study of patients with brain lesions and well-defined symptoms. Such studies required patience, as individual brain anatomy could only be revealed after death. So research projects might well last a lifetime for some or remain unachieved for others. Nevertheless, studies of single cases or case series revealed the first white matter biomarkers for neurological syndromes such as conduction aphasia ^6^, transcortical motor aphasia, subcortical motor aphasia, transcortical sensory aphasia, subcortical sensory aphasia ^7^, alexia without agraphia ^8^, bilateral and unilateral apraxia ^9^. These historical contributions demonstrated that the proper **functioning of the brain** can be disrupted even by distant lesions through disconnection and diaschisis mechanisms ^10,11^.

Fundamental advances in our understanding of brain function came from this work. Nevertheless this static vision of the brain did not account for classical conditioning effects ^12^ or memory ^13^, or diaschisis ^14–16^ phenomena that required notions of **plasticity**. Donald Hebb introduced this concept as follows “*Let us assume that the persistence or repetition of a reverberatory activity (or “trace”) tends to induce lasting cellular changes that add to its stability.… When an axon of cell A is near enough to excite a cell B and repeatedly or persistently takes part in firing it, some growth process or metabolic change takes place in one or both cells such that A’s efficiency, as one of the cells firing B, is increased.*” With these words, Hebbian theory acknowledged the possibility for changes to occur in the brain, and provided a framework for conceptualising learning through the mechanisms of plasticity ^17^. However, measures of **plasticity** were quite challenging, as they required the anatomical investigation of living samples with very invasive methods.

Decades later, magnetic resonance imaging (MRI) brought a new window on human brain anatomy. In 1985, a breakthrough came with the emergence diffusion-weighted MRI (DWI, ^18^). DWI measures the diffusion of water molecules along different directions. Given that axons are impermeable they constrain water diffusion to their main direction. Hence, DWI indirectly assesses white matter microstructure. Further by piecing together local estimates of water diffusion orientation, main bundles of axons can be reconstructed. This technique clearly boosted research because of its non-invasiveness. **Anatomical studies** performed in living human brains were able to refine the description of large fiber bundles ^19,20^, allowing for replication in large numbers of participants ^21,22^ and enabling the discovery of inter-individual variability ^23,24^. **Functional models** could now benefit from whole brain connectomes for the first time ^25^, identifying new disconnection syndromes ^26^ and allowing for new divisions of the cortex based on axonal inputs and outputs or connectivity based parcellation ^27,28^. Finally, the structural connectome of an individual could be assessed different time points allowing for the investigation of white matter **plasticity** for the first time ^29^.

Hence, in this review of the role of DWI in neuroscience we will first describe the anatomical advances provided by this method and will then survey its importance in the establishment of new brain models. Finally, we will review studies that investigated white matter plasticity. Hopefully, the ideas expressed in this review will provide a comprehensive understanding of the scientific question investigated with DWI and encourage new generations to pursue exploit this powerful technology to gain novel insights into human brain wiring.

## Part 1/ Anatomy

Diffusion MRI provides different types of information about human brain anatomy. Local (voxel-wise) measures of diffusion properties can give insight into local white matter microstructure, whereas exploiting these measures to perform in-vivo tractography can provide information on the organisation of white matter at a systems level.

### Local measures of white matter microstructure

Voxel-wise measures of diffusion properties, such as fractional anisotropy or mean diffusivity, are modulated by local tissue microstructure ^30^. For example, studies in model systems show that fractional anisotropy increases with increasing packing density or decreasing axon diameter ^31^, and decreases with reductions in myelin ^32^. However, although empirical or simulation studies can clearly demonstrate that varying these tissue properties modulates diffusion parameters, we are still faced with an inverse problem when trying to interpret an observed difference in diffusion parameters (e.g. reduced fractional anisotropy), as there is not a one to one relationship between a given diffusion parameter and the underlying tissue structure ^33–37^. Nevertheless, diffusion parameters provide useful non-invasive measures of local tissue microstructure, with sensitivity to features including membrane integrity, myelin thickness, axon diameter and packing density. These physical characteristics of the white matter fibre bundle will have consequences for the physiological functioning of that bundle, affecting properties such as conduction time, refractory time, probability of transmission or even synchronisation of signals across a distributed cortico-cortical network. Variations in these physiological properties may in turn be expected to give rise to variation in behavioural outputs ^33,38,39^.

For instance, left and right hemispheres show variation in function as well as in anatomy. Diffusion parameters’ comparison between the two hemispheres indicated increased fractional anisotropy in the left hemisphere in the external capsule ^40^, cingulum bundle ^41^ and perisylvian white matter ^42^ thought to support left hemispheric dominance for language functions ^43,44^. Right asymmetries were also identified in the dorsal fronto-parietal white matter ^42^ that could be related to the right hemispheric dominance for spatial processing of information. However, the high variability between subject and the lack of behavioural measures in these preliminary studies did not allow a solid relationship to be drawn between voxelwise measure of interhemispheric differences and functional dominance. Particularly, the relationship between language lateralisation and structural connectivity measures in the language system remains intricate ^43,45^.

Further studies have shown that individual differences in measures of local white matter microstructure correlate with variations in physiological properties of the fiber pathways ^46–52^. For example, paired-pulse transcranial magnetic stimulation can be used to probe the functional connectivity of a cortico-cortical connection. A conditioning pulse applied to dorsal premotor cortex of one hemisphere will modulate the excitability of the primary motor cortex in the other hemisphere; the degree of modulation can be used to estimate functional connectivity between the two cortical areas. Individual differences in this measure were found to correlate with variation in white matter fractional anisotropy (FA), such that individuals with stronger functional connectivity had higher FA in callosal or subcortical white matter pathways between premotor and motor cortex ^53^. This supports the idea that variation in white matter microstructure is associated with variation in the physiological properties of those fiber bundles.

Advanced diffusion weighting imaging analyses provide estimates of the axon diameter distribution or fibre composition along with other physical properties, such as the intra-axonal resistance, membrane resistance and capacitance, etc, help determine many important functional properties of nerves, such as their conduction velocity or rate of information transfer ^54–58^. Neural pathways that are characterized by fast reaction times (e.g. within the motor system) will also exhibit higher percentage of large-diameter axons. On the other hand, pathways that show slow conduction velocity will also exhibit a larger population of smaller axons or even non-myelinated axons. Therefore, it is only logical to assume that the properties of axon diameter distribution will have critical role on healthy central nervous system functioning and will be dramatically affected in abnormal conditions and disease.

Despite the fundamental new insights it provides, diffusion weighted imaging has several important limitations. The key limitation is implicit in the diffusion tensor model (DTI, ^59^). This assumption oversimplifies the true motion of water within brain tissue. The practical consequence of this over-simplicity of the diffusion tensor model is that the common DTI indices, mean diffusivity and FA, are non-specific to any particular tissue compartment. Features of microstructure, such as cell size, density, permeability and orientation distribution, all affect DTI indices and changes in the indices are impossible to associate with more specific changes in microstructural features. Recent trends aim to use more sophisticated models of diffusion in order to measure microstructural features directly ^60,61^.

One such approach for modelling the diffusion signal (instead of DTI) is to devise a mathematical formalism that is guided by tissue geometry. Yet, anyone who has looked down a microscope at a brain section would have been impressed by the complexity of the white matter geometry ^36,62^. Cells of different shapes and sizes with processes spreading at different scales and orientations depict a mess of geometries at the micron-scale. Yet even at a lower-level of magnification, where cell layers and fibre bundles are visible, complexity arises where fibres cross, disperse and fan within hundreds of microns.

Pioneering work by Stanisz and collaborators attempted to reduce the complexity of neural tissue modelling by introducing a two-compartment geometrical framework that includes diffusion within spheres and cylinders ^63^. Further works have suggested that diffusion within the confined boundaries of cells and axons can be regarded as restricted diffusion. Under this approach it is hypothesized that the geometry of the tissue affects the diffusivity of water molecules. For example, it is reasonable to assume that water diffusion within the axon will be restricted by the myelin membrane while elsewhere it will be only hindered or free. This is the basis for the composite hindered and restricted model of diffusion (CHARMED) ^60,64^. In CHARMED, restricted diffusion in the intra-axonal space is modelled as diffusion within impermeable cylinders ^64–66^. As a consequence, the CHARMED model allows enhanced characterization of the axonal water compartment. One of the advantages of CHARMED is that it is possible to reconstruct the 3D displacement distribution function for each of the components (hindered and restricted). Thus, it is possible to extract physically meaningful parameters for each of the diffusing components; these parameters include the diffusivity of the extra-axonal matrix (diffusivity of the hindered part), the axonal density (the volume fraction of the restricted part), and the fibres directions (the orientation density function of the restricted part). While the CHARMED model provides a conceptual framework to separate different modes of diffusion and relate them to tissue compartments, it allows modelling of 2-3 different fibre orientations with accuracy limited by the number of measured directions ^60^. Subsequent works tried to expand the CHARMED model to include fibre dispersion and fanning within a voxel and increase the accuracy and orientation estimation of the model ^67–71^. Noteworthy is the NODDI framework which expands CHARMED to model also dispersion of fibre orientation but with optimized acquisition and analysis pipelines ^69^.

Neural tissue is complex, and it should be realized that there will not be a single model that can capture all the details and complexity of different neural compartments ^62,72^. The different abovementioned models (and others) try to separate different features of interest of the tissue. Obviously including too many free parameters in the model will lead to computational problems such as over fitting and other optimization and data fitting issues. Thus, the optimal model for diffusion imaging depends on the research question. For example, an important feature of white matter that is not modelled in CHARMED or NODDI is axon diameter ^73–75^. Being an important feature of brain connectivity with implications for conduction velocity and information transfer efficiency within the brain ^76,77^, the in-vivo measurement of axonal diameter became another modelling challenge. In an extension of the CHARMED framework, called AxCaliber, the estimation of the axon diameter distribution became also feasible ^78,79^. The idea behind AxCaliber is that each axon size will experience restricted diffusion at a different diffusion time. For example, an axon with a diameter of 1um will experience restricted diffusion already at very short diffusion times while a larger axon will experience restricted diffusion only when the diffusion time is increased. By acquiring a multi-diffusion time CHARMED data set it is possible to accurately estimate the axon diameter distribution function with the AxCaliber framework. In AxCaliber, unlike CHARMED, water diffusion is measured exactly perpendicular to the long axis of the fibers (this is a prerequisite of the model). It was found that by using a gamma function (with only 2 free parameters) it is possible to adequately estimate the axon diameter distribution function by AxCaliber ^79^.

The AxCaliber framework was verified on excised samples of optic and sciatic nerves. These two nerve samples have very different axon diameter distribution functions (**Figure 2**). The AxCaliber diameter distribution functions were in good agreement with the histological ones (**Figure 2**) ^79^. In addition, AxCaliber was implemented to study the morphology of the corpus callosum (in-vivo). AxCaliber analysis was performed on a voxel-by-voxel basis providing for each the axon diameter distribution function for each voxel. These distributions were used as an input to a clustering algorithm in order to visualize regions with significantly different axon diameter distribution (ADD). The clustering based on the ADD was able to segment the corpus callosum into several regions that fit known morphological zones of those samples ^80^ (**Figure 3**). Recently the axon diameter distribution was demonstrated in the human brain and also was shown to correlate with conduction velocity measures of interhemispheric transmission time ^81^. Hence correlations between in vivo measures of axonal diameters and reaction time measures during cognitive paradigms will be of particular interest in near future.

**Figure 2:**
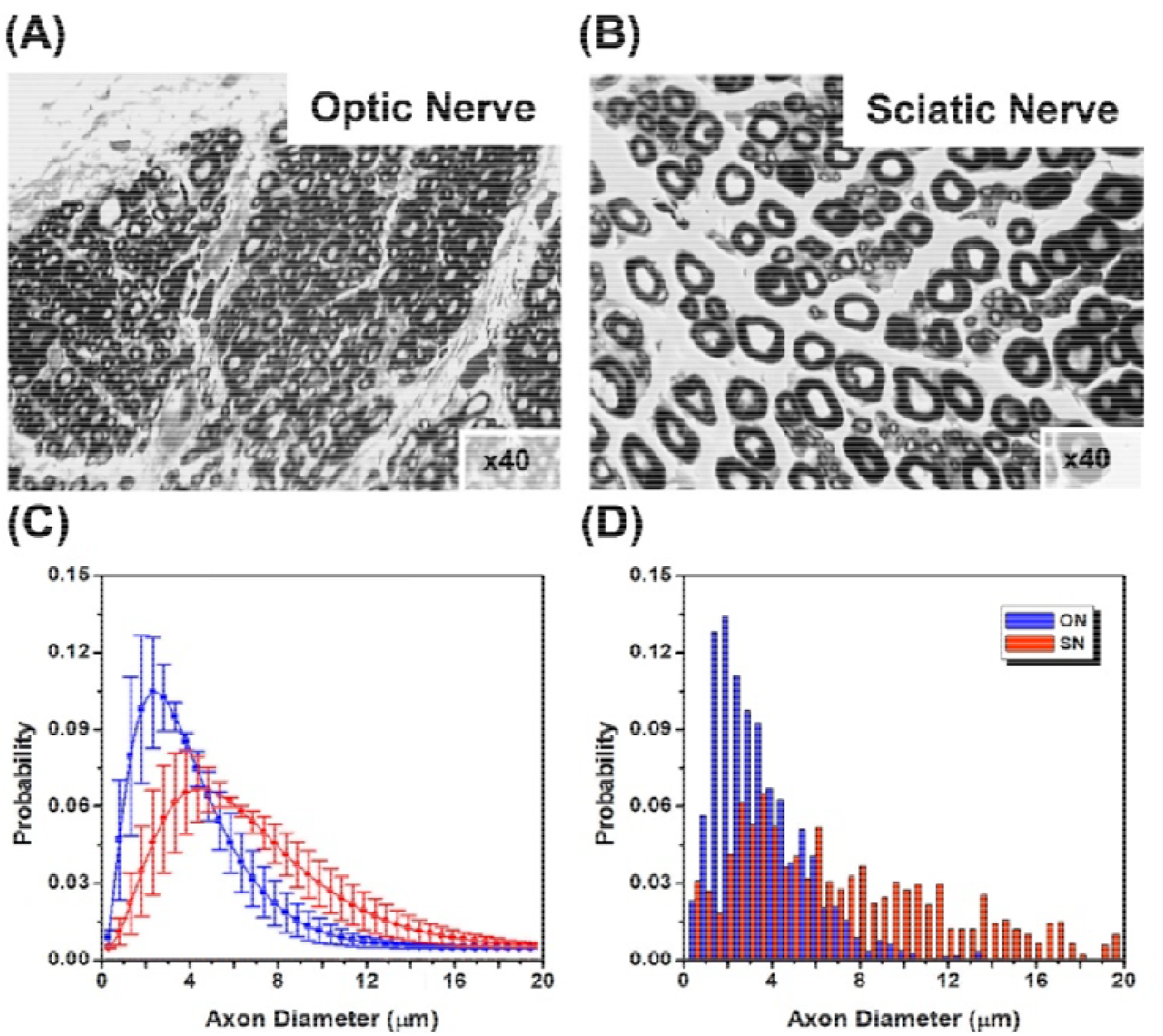
AxCaliber of porcine optic and sciatic nerves: (A) and (B) are electron microscope section of optic nerve (A) and sciatic nerve (B). (C) AxCaliber axon diameter distribution - (optic – blue, sciatic –red). (D) Axon diameter distribution derived from electron microscopy

It should be noted that the original AxCaliber framework suffers from several inherent limitations. The first is the need to measure the diffusion properties exactly perpendicular to the fibre orientation. This limited the application of the method to the corpus callosum. In a further development of the CHARMED and NODDI frameworks it was suggested that the mean axon diameter could be directly measured for any orientated fibre system ^82^. The ActiveAx acquisition and analysis pipeline, although not measuring the entire axonal distribution probability function, provides an approach to estimate the mean axon diameter for the entire brain. Another limitation to measuring axonal diameter properties with diffusion imaging is the need to have high diffusion gradient amplitude in order to have comparable accuracy over a wide spread of axon diameters. It was shown, theoretically, that small axons (<1micron) will be inaccurately estimated with current gradient technology and will bias the overall measured axon diameter ^83,84^. This measurement artefact is significantly minimized with new gradient technology providing high amplitudes of diffusion gradients (300mT/m) ^85^. Yet, other works suggested that the modelling algorithm can be further developed to cope with the abovementioned limitations by introducing additional features such as restricted diffusion within the extracellular space ^86–88^. Yet, despite this limitation, the measurement of axonal properties provides an additional micro-structural feature that can better characterize brain connections and connectivity.

**Figure 3:**
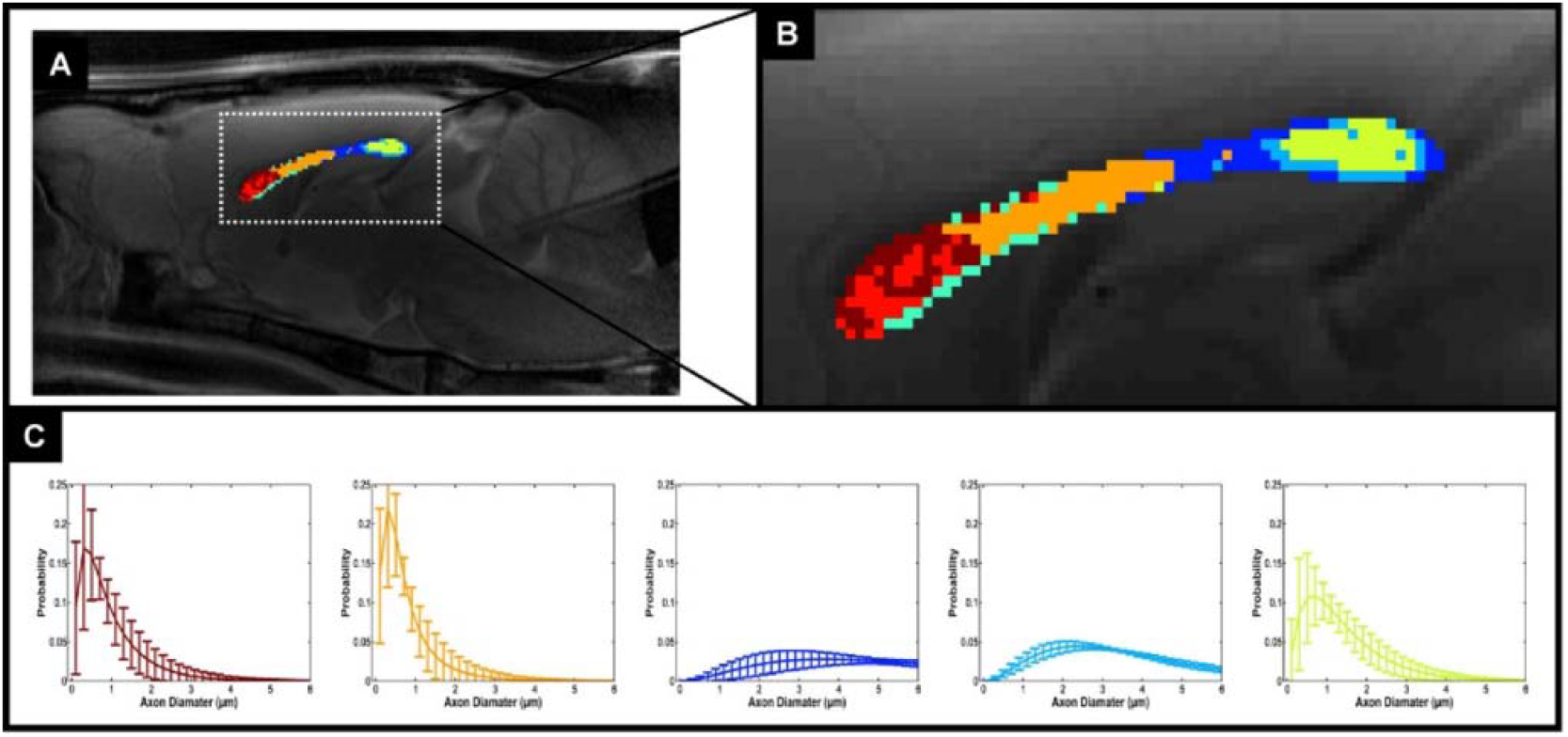
Cluster analysis of Ax-Caliber’s ADD along the corpus callosum: (A) A mid-sagittal T2-weighted MRI with the AxCaliber clusters super-imposed, enlarged at (B). (C) The AxCaliber averaged ADDs for the different clusters given in (A) and (B).

### Tractography to estimate long-range connectivity

As mentioned above, local estimates of dominant diffusion directions can be followed to reconstruct estimates of fiber pathways. There are many different methodological approaches to performing diffusion tractography, as reviewed elsewhere in this special issue ^37,89^. From the neuroanatomical perspective, tractography allows us to estimate the organisation of major fibre pathways in the human brain (**Figure 4**) and to explore variation in the strength and organisation of these pathways between individuals or over time.

**Figure 4:**
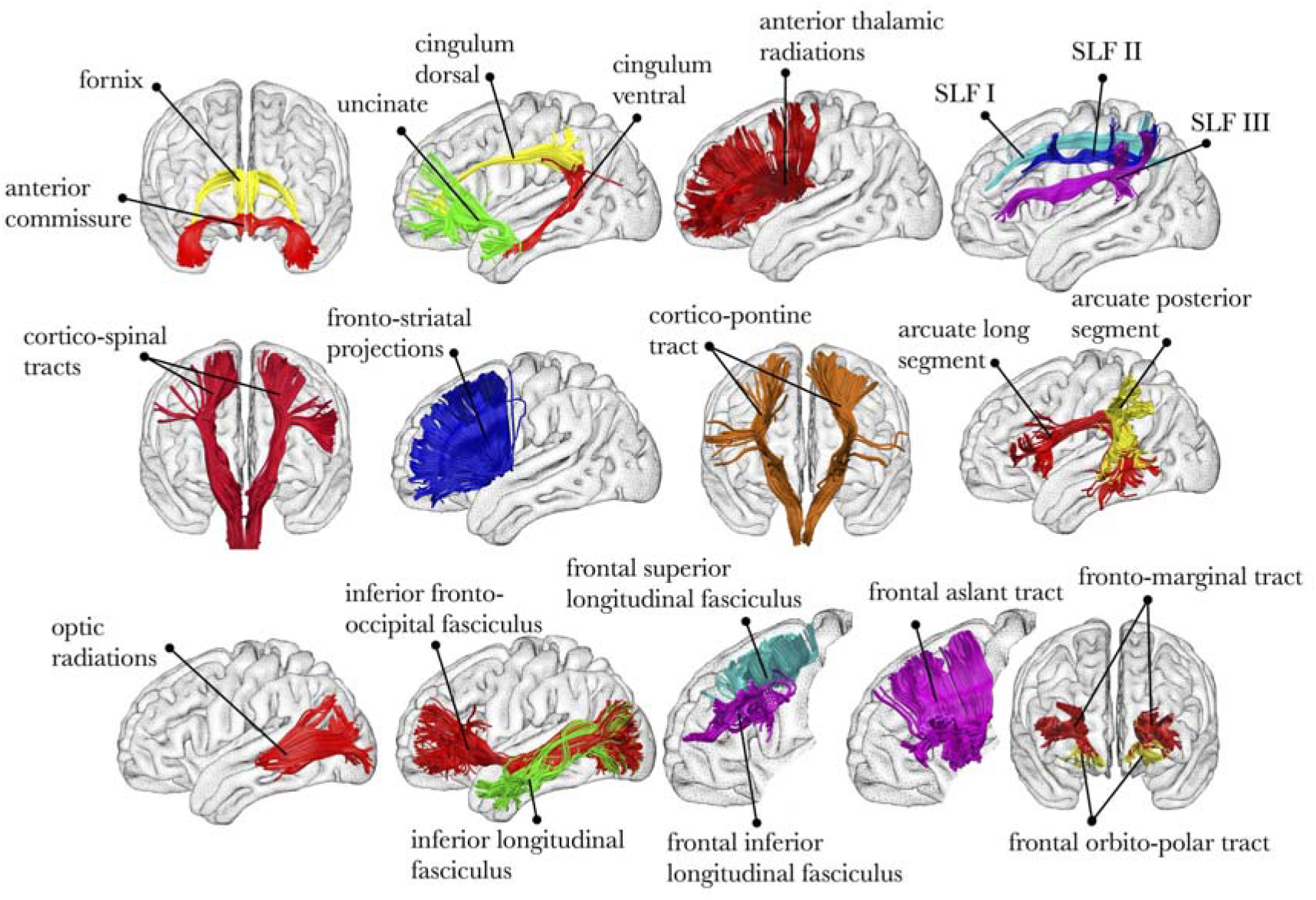
Virtual dissection of the major fibre pathways in the human brain ^90^.

Early applications of diffusion tractography were able to provide the first ‘in vivo dissections’ and mapping of major white matter fiber bundles in the human brain with an increasing precision ^19,21,91–100^ and led to the publication of several atlases ^101–103^.

The use of tractography for anatomical studies has mostly depicted associative tracts (i.e. cortico-cortical connection) as they were defined in the early XIXth century (**Figure 5a**). Handling of tractography was indeed much easier than standard post mortem white matter dissections and led to the discovery of new associative tracts which were consequently replicated using standard post-mortem Klingler dissections ^104,105^. In that sense tractography has boosted our anatomical knowledge of white matter pathways by providing an easier access to white matter connections ^106,107^. For instance, much evidences from tractography, cross validated with post-mortem dissections, now converges toward a model of the arcuate fasciculus splitted into three branches (temporo-parietal, fronto-parietal and posterior parietotemporal, **Figure 5b**). However, caution is required to be taken as many limitations in tractography may mislead neuroanatomists by piecing together different tracts segments into large single bundles ^37,89^.

**Figure 5:**
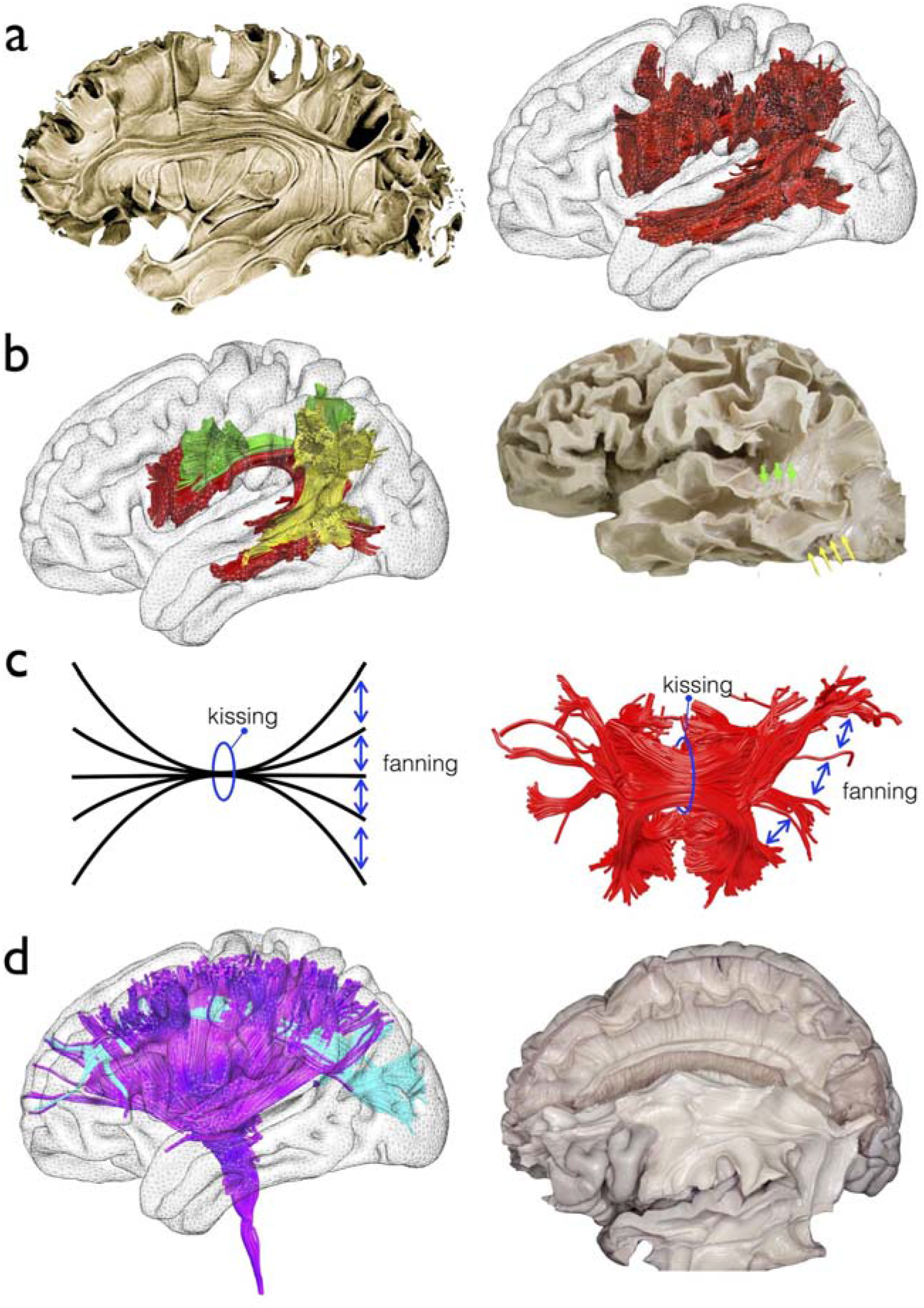
White matter anatomy from early tractography to advanced models. a) Standard anatomical description of the arcuate fasciculus (Left panel, ^104^) replicated with diffusion weighted imaging tractography (Right panel, ^19^) b) New anatomical model of the arcuate fasciculus (Left panel, ^20^) validated with Klingler post-mortem dissection (Right panel, ^95^) c) Model of white matter configurations that challenge classical tractography (Left panel), example of the tractography of the corpus callosum splenium showing the same limitation (Right panel). d) First branch of the superior longitudinal fasciculus crossing with the corona radiata discovered in humans with advanced spherical deconvolution tractography (Left panel, ^77^) later validated with Klingler post-mortem dissection (Right panel, ^117^)

This is particularly the case for projection (i.e. subcortico-cortical) and commisural pathways (i.e. interhemispheric), which have been more challenging to dissect due to severe limitations in the tractography of complex white matter configurations (i.e crossing, fanning and kissing configuration) (**Figure 5c**). Although crossing has been the focus of interest of much work for the past 5 years, with very promising results particularly evident with the use of spherical deconvolution tractography ^108–113^ (**Figure 5d**), fanning and kissing configurations still remain a clear challenge in this field of research. The fanning limitation is characterised by an underestimation of the fibre count reaching an area and the kissing limitation by the misidentification of the correct group of fibres to track. Together, these two problems hamper progress in the depiction of the projection and commissural pathways ^114–116^.

### Tractography to divide grey matter into subregions

At a cortical level, SMA and pre SMA, two areas working hierarchically in the control of action ^118^ have a different pattern of connection with the rest of the brain. Exploiting this difference, connectivity information gained from diffusion tractography can be used to define the boundary between these two regions based purely on connectivity, driven by the fact that SMA shows a closer relationship with primary motor cortex whereas pre-SMA is more connected to the prefrontal cortex ^27^. Indeed, connectivity-based parcellation has been used to define subdivisions across a range of different human cortical areas ^119–126^, as well as to demonstrate homologies between human and monkey cortex ^127–133^. Recently, diffusion tractography was used along with other multimodal structural and functional MRI data to generate a novel whole brain cortical parcellation using data from the Human Connectome Project (**Figure 7a**, ^126^).

Alongside enthusiasm for the newfound ability to visualise white matter bundles in living human brains, have been notes of caution to keep in mind that tractography is not the same as invasive tract tracing. ‘Gold standard’ invasive tract tracing typically involves injecting a tracer into an area of interest, allowing it to be taken up by cells, and then observing it in interconnected regions after it has been transported there via anterograde and/or retrograde transport mechanisms. As such, invasive tract tracing involves tracing axonal pathways. By contrast, in diffusion tractography we are typically following estimates of the path of least resistance to diffusion. In many cases, this corresponds roughly to the path of the dominant fibre orientation, but in other cases it does not. There is therefore awareness that, as with any method in biology, diffusion tractography is susceptible to false negatives (inability to track pathways that we know exist) and false positives (tracking spurious pathways). Much valuable work has been done to try to validate diffusion tractography, giving us a clearer idea of the limitations and strengths of the method ^134–141^. This is challenging work, requiring development of novel phantoms with realistic properties ^115,142^, or meticulous reconstruction of fibre pathways from brain sections taken from tract tracing experiments ^143,144^. Such work is encouraging but there is more to be done to allow neuroscientists to be able to interpret diffusion tractography results with confidence ^145^.

### Part 2/ Functional models

Functional models of the brain originally emerged from the study of brain damaged patients ^146^. The localisation of the lesion in the brain was indeed interpreted as the core origin of the functional impairment ^147^. However, the localisation of the lesion appeared insufficient to explain the existence of patients with different lesions and similar symptoms. The idea emerged that the communication between these regions was impaired through a mechanism of white matter disconnection. Traditional model of disconnection syndromes includes conduction aphasia ^6^, transcortical motor aphasia, subcortical motor aphasia, transcortical sensory aphasia, subcortical sensory aphasia ^7^, alexia without agraphia ^8^, bilateral and unilateral apraxia ^9^. With the advent brain imaging the neuroimaging community focused on voxels independently ^148–150^ without any attempts to capture correlation across them ^151,152^, effectively masking disconnection syndromes ^153,154^. The access to visualisation and mapping of white matter connections instead allowed for the investigation of white matter connections in patients and led to the discovery of new disconnection syndromes. For instance, visuospatial neglect, a severe neurological condition characterised by the loss of half of the visual field’s awareness, is associated with the disruption of the fibres connecting the frontal to the parietal lobe ^26,153,155^. Tractography also contributed to the establishment or confirmation of a new disconnection hypothesis for Gerstmann syndrome ^156^, and neurodevelopmental hypoconnectivity hypotheses for psychopathy ^157^, dyslexia ^158^ as well as congenital prosopagnosia ^159^. Conversely, aberrant/increased connections (i.e. hyperconnection) hypotheses have recently been suggested as a potential mechanisms for inattention in attention deficit hyperactive disorders ^160^, auditory ^161–163^ and visual hallucinations ^164^ in schizophrenia. All these results demonstrate the utmost importance of brain connection to the proper functioning of the brain. Indeed brain areas deprived of their inputs (i.e. afferent connections) or outputs (i.e. efferent connections) will no longer be able to contribute anymore to the elaboration of the cognition and the behaviour.

With regards to effects on behaviour, studies have shown that individual differences in behavioural performance for a given task correlate with variation in white matter microstructure of task-relevant pathways, even in young healthy populations ^39,165 38^. For example, as summarised in **Figure 7**, variation in bimanual co-ordination skill correlates with variation in fractional anisotropy in the body of the corpus callosum, a white matter area that contains transcallosal pathways between primary and supplementary motor areas ^166^. Similar effects have been found in a wide range of sensory, motor and cognitive domains including for example vision ^167–169^, audition ^170^, motor skills ^171–175^ and language ^176,177^, literacy ^178–180^, emotion ^181^ and motivation ^182^, visuospatial ^183^, memory ^184^ and executive function ^175,185,186^ and individual characteristics such as creativity ^187^, musical skills ^188^ and personality ^189,190^.

Such evidences suggest that the function of a brain area can be defined by its afferent connections from and efferent connection to other areas ^192–194^. As reported in the previous section, brain regions can be characterised by their structural connectivity with the rest of the brain. Therefore, boundaries based on structural connectivity might well give a functional parcellation of the brain. Preliminary evidence starts to abound in this direction. For example, delineating subregions of human thalamus is important to be able to study the function of these regions in vivo and to be able to target specific nuclei for surgical interventions such as deep brain stimulation for movement disorders. However, divisions between thalamic nuclei are not visible on standard structural MRI. Using diffusion tractography, thalamic subregions can be reliably identified based on cortical connectivity patterns ^28^. These connectivity-based divisions have been shown to have functional relevance ^195,196^, and could be useful for individualised neurosurgical planning ^197^. Recently, tractography revealed a division of the frontal lobes in 12 areas characterised by their connection with the rest of the brain and a clear-cut functional specificity (**Figure 5b**)^126^. Hence model of structural connectivity assessed with tractography seems to capture the functional organisation of the brain.

However, the past two decades have been marked by the discovery of large interindividual variability in brain structure and function (i.e. different phenotype), creating additional challenges in furthering the understanding of the brain and a lack of models to explain its functioning ^198–201^. Studies have begun to explore and describe these pathways across large numbers of individuals ^24,116,202^. Just as studies have found that individual differences in behaviour correlate with variation in local white matter microstructure, so too have tractography studies found that variation in the ‘strength’ or organization of white matter fibres relate to variations in behaviour. For instance, language areas and their interconnections are predominantly represented in the left hemisphere for most but not all healthy participants ^203,204^. However, individuals vary in the degree of lateralisation and people with more symmetric patterns of connection have been shown to perform better at episodic memory tasks ^203^. These differences are important consideration for the field of clinical neuroscience. Forkel et al. ^205^ demonstrated that the phenotype of the structural network supporting language (i.e. arcuate fasciculus) may interact with recovery after a stroke in the left hemisphere. Additionally, Lunven et al. ^206^ reported that the strength of inter-hemispheric communication is important for the recovery of visuospatial neglect after a stroke in the right hemisphere (see ^207,208^ for extensive reviews on the role of diffusion MRI in clinical neurosciences). Hence the study of inter-individual variability is a new line of research directly accessible in large scale projects such as the Human Connectome Project (https://www.humanconnectome.org), where high quality diffusion MRI data have been acquired in 1200 subjects and openly shared with the community ^209^. Using diffusion weighted imaging tractography to characterise the functional specialisation of brain areas at the individual level may therefore benefit the domain of inter-individual variability, providing a more tailored brain model when doing group comparisons eventually making methods like spatial registration and smoothing obsolete.

## Part 3/ Measures of Plasticity

Until the mid ’90s, research on white matter was limited to histological investigations of its anatomy, for two reasons: 1) There was no in-vivo and non-invasive imaging method that could provide quantitative measures of the white matter; 2) it was thought that in the fully developed and healthy brain, the micro-structure of white matter should be roughly stable and fixed. Nevertheless, recent studies suggest that the white matter can undergo changes following training or cognitive experience. Such studies coined the term white matter plasticity.

Neuroplasticity, the functional and structural re-organization capacity of the brain may occur at several levels (from molecular to regional changes) and time-scales (seconds ^210,211^ to years ^212^). Obviously, the definition of neuroplasticity and its investigation are centred around neurons and less on white matter. Nevertheless, the scope of neuroplasticity is wide, ranging from short-term changes at the synapse through long-term potentiation (LTP) and depression (LTD) mechanisms through long-lasting new neuronal connections (dendritic trees) ^213–218^.

Pioneering studies revealed that new white matter projections can be formed following extensive learning procedures ^219,220^. These studies, performed on post-mortem monkey brains using tracer methods revealed that the brain can rewire itself concomitantly to cortical neuroplasticity ^221^. Recent works utilized diffusion tensor imaging to reveal white matter changes following long term training (weeks to months) ^180,188,222^. These studies suggested that diffusion imaging can be sensitive to white matter plasticity. Recent evidences showed that following only a 5-day water maze task, in addition to significant cortical plasticity, white matter plasticity in the corpus callosum could be detected with DTI ^223^. Histology showed that this plasticity, characterized by increase in FA, is also manifested by elevated levels of myelin (Figure 6). In addition, a recent molecular imaging study suggests that already following action potentials, signalling mechanisms control oligodendrocyte myelin formation 224. This experiment was conducted on cultured neural tissue. Indeed, a recent study showed that 2 hours of spatial navigation task are sufficient to cause diffusion changes in the fornix. All these studies on plasticity are not exempt of limitation ^225^ and future research is needed in order to strengthen that field.

**Figure 6:**
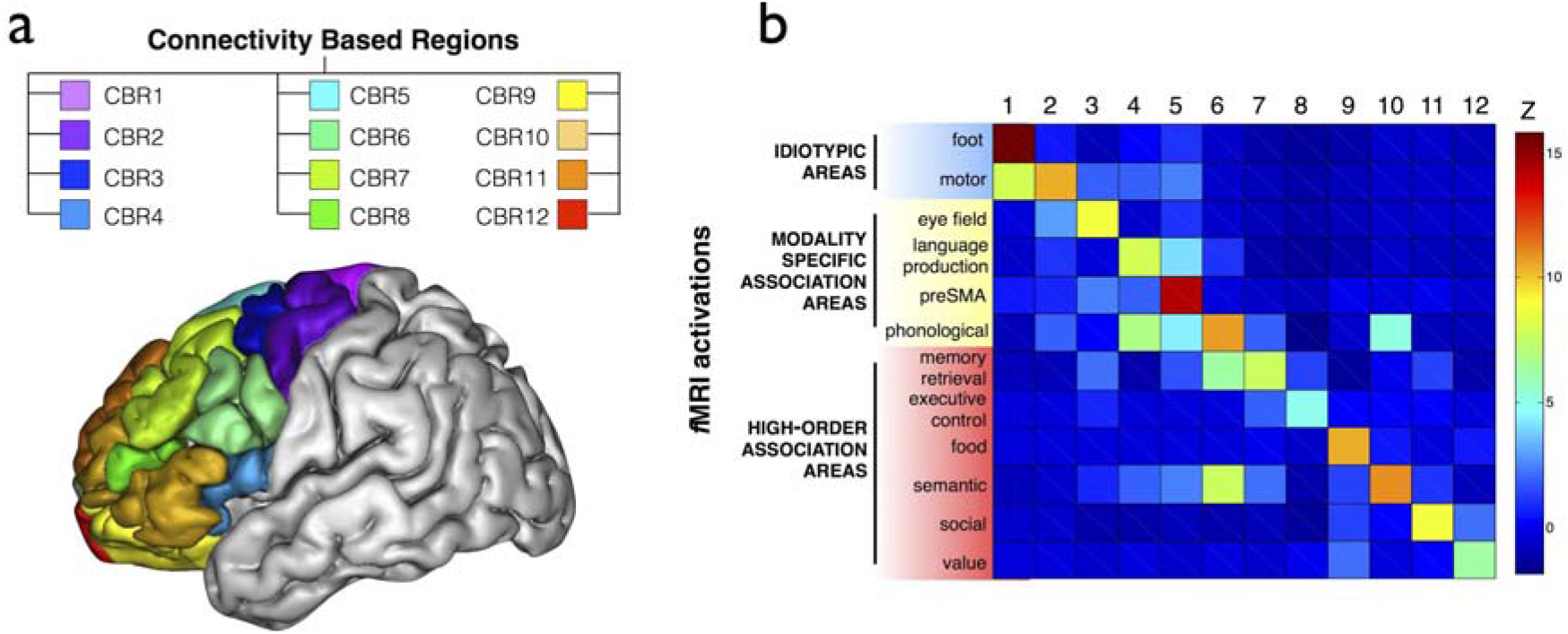
Connectivity-based parcellation of the frontal lobes. a) 3D lateral view of the brain and the 12 regions defined by the structural connections with the rest of the brain. b) functional specificity of the 12 connectivity based regions.

**Figure 7:**
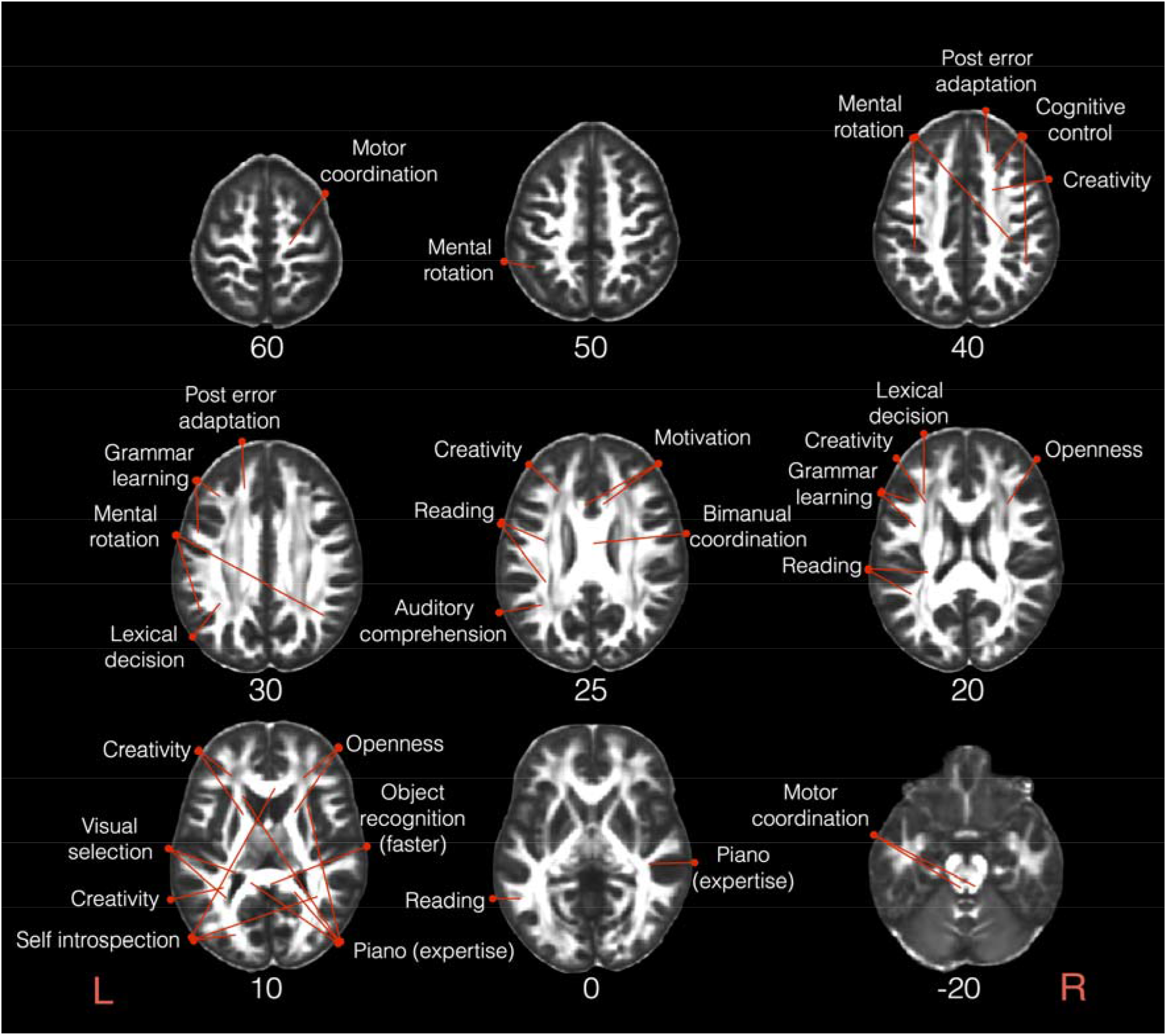
Correlations between individual differences in behavioural performance for a given task and variation in white matter microstructure. Axial slices are displayed in neurological convention. Numbers indicate MNI152 axial coordinates ^191^. L, Left; R, Right.

**Figure 8:**
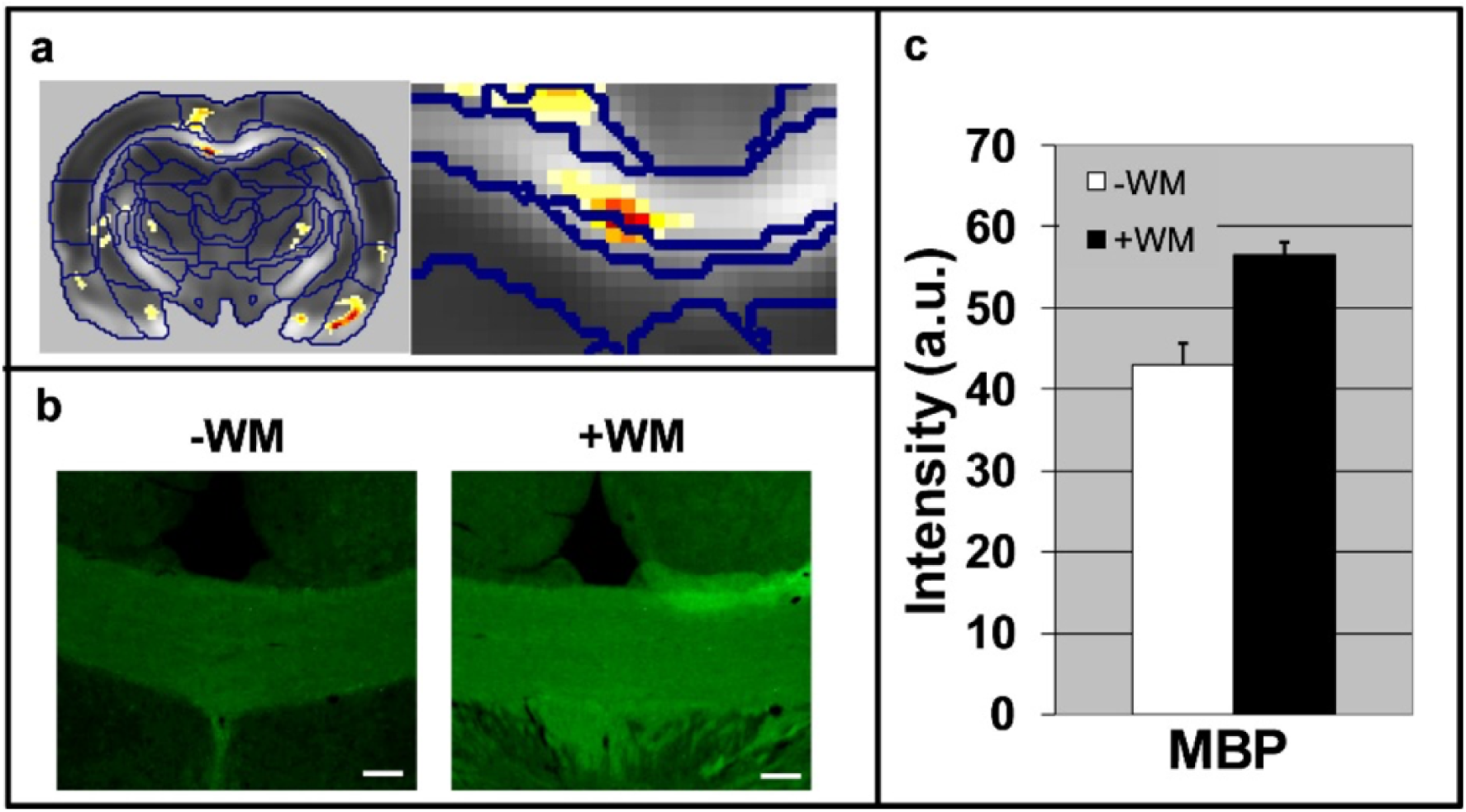
White matter plasticity following a 5-day water maze. (a) Statistical parametric maps of interaction in FA values between scan time (pre vs. post water maze) and group (learning vs. control). An enlargement of the corpus callosum region is shown on the right. (b) Immunohistochemical staining (x10 magnification) of the CC for MBP. Note the increase in MBP immuno-reactivity in the CC after the water maze task. (c) Quantification of the immunoreactivity (staining intensity) in the CC (P< 0.05).

These experiments, although performed on different species and samples (human, rodents and tissue culture) suggest that the white matter is a dynamic tissue. Yet, what exactly happens in the white matter is not fully understood and the temporal evolution of this phenomenon is uncharted. Specifically, a significant impact on neuroscience will be the demonstration of white matter plasticity in the human brain following short episodes (minutes/hours) of training. We anticipate that short-term white matter plasticity will be stimulate different mechanism from long term plasticity, however, DTI might be not sensitive and specific enough to demonstrate that difference. More specific methods such as CHARMED and AxCaliber could provide the enhanced sensitivity and specificity needed for comprehensive characterization of this phenomenon. Nevertheless, once white matter plasticity mechanisms can be resolved with MRI, these kinds of experiments could shed a new light on brain physiology and provide a whole new definition to the concept of brain connectivity.

## Conclusion

Over the past 20 years, diffusion weighted imaging has proven to be an indispensable approach for the study of white matter. It has provided invaluable insight into neuroanatomy, which has led to the discovery of new pathways, and refinement of what we already knew. Notably, diffusion weighted imaging has demonstrated the existence of a clear organisation and a true inter-individual variability in the way that the brain is connected, and that this variability directly impacts the functioning of the brain. Finally, diffusion imaging offered the unique opportunity to directly quantify brain plasticity. However, while reducing the gap between direct biological and indirect DWI quantification of brain tissues continues to be a challenge, it is a necessity for future generations who may wish to exploit this powerful technology to gain novel insights.

## Acknowledgements

We wish to thank Lauren Sakuma for useful discussion. This work was supported by the *‘Agence Nationale de la Recherche’* [grants number ANR-10-IAIHU-06 and number ANR-13-JSV4-0001-01].

